# The Spontaneous Evolution of Biology

**DOI:** 10.64898/2026.07.14.738476

**Authors:** Sara ten Have, Ewan McMillan, Tim Medway, Rob Kent, Alan Prescott

## Abstract

We have demonstrated the potential of amino acids to polymerise into peptides, proteins, and form cell-like structures in the absence of cellular machinery, including nucleic acids, lipids or sugars. Not only has cell-free protein replication been observed, but evidence of protein templating strongly suggests protein mediated replication. We believe this is the first experimental demonstration of the link between the Miller-Urey experiment which produced amino acids from elemental starting material, and cell-like structures.

Life, by definition, is the condition that distinguishes animals and plants from inorganic matter, including the capacity for growth, reproduction, functional activity, and continual change preceding death. Here we have characterised peptides which form reproducibly, into structures with longevity and which subsequently catalyse the polymerisation of free amino acids into copies of themselves. The proteomic analysis of these samples over time also enables evolution of peptide sequences to be seen and quantified. This evolution of both complex structure and functionally active proteins may potentially demonstrate a credible path to the beginnings of life, which we call the Spontaneous Evolution of Biology (SEB) Theory.

**One Sentence Summary:** We have shown how peptides and proteins can be made from amino acids in an aqueous media without cells, lipids or nucleic acids, duplicating themselves and forming complex structures which resemble cells.

## Introduction

The estimated environment from which life arose 4.6 billion years ago defines that water was essential for life (*1-5*). Further, the experiments conducted by Millar and Urey(*6-8*) demonstrate, from the likely conditions of early earth, the formation of amino acids and more complex carbon-based molecules derived from gases, water and energy sources. There remains a gap in the knowledge from amino acid formation to polymerisation into peptides and proteins. Polymerisation in simple conditions is being described by discoveries showing peptide bond formation in water droplets, from single amino acids (*9, 10*), and templated peptide synthesis from peptides halves, containing an activated thioester and a cysteine(*11-14*). These recent discoveries show that amide bond formation in water is possible and frequent without complicated chemical modification, enzymes or other catalysts, despite the energy required for these bonds to form. Additionally, they demonstrate the templating property has a catalysing effect on peptide fragment annealing, verifying information transfer from peptide to peptide and showing templating capabilities for peptides.

The mechanism for the first generation of complex life, from the building blocks-DNA, RNA and amino acids - has not been characterized previously. Several theories for how this may have come about have been hypothesized (such as RNA World (*15*), Panspermia (*16*), and Urzyme (*17*) theories) but not experimentally proven. For example, RNA world relies not on RNA as it’s modern-day form being the precursor but simpler RNA-like polymers being the progenitors but “*We do not have any remnants of these compounds in present-day cells, nor do such compounds leave fossil records. Nonetheless, the relative simplicity of these “RNA-like polymers” make them better candidates than RNA itself for the first biopolymers on Earth”*(*18*). More recently RNA enzymes have been constructed which enabled replication and evolution(*19*)but with highly engineered components and traditional PCR processes-which is a significant step from nucleic acids in insolation. Urzymes additionally had been hypothesized and artificially synthesized to test their minimal enzymatic capabilities, but recently there have been naturally occurring minimal enzymes which are similar to the hypothesized counterparts(*20*) Panspermia, while an interesting prospect, does shift the progenitors for life off planet, and therefore conveniently shifts reasoning and the burden of proof “elsewhere”. The central molecular focus is DNA and RNA, due to the perception that these molecules are the basis of, and control Life. The Spontaneous Evolution of Biology (SEB) theory, and its experimental evidence, suggests that amino acids and the resulting peptides and proteins are capable of performing the core facets of life (the capacity for growth, reproduction, structure, functional activity, and continual change preceding death(*21*)) independently of DNA and RNA, or cellular machinery.

The implication is that protein formation independent of DNA/RNA is plausible, given the wide ranging essential biological roles of proteins. The cell structure is made from protein, signals are sent via proteins and peptides(*22*), and transfer in and out of the cell is mediated by protein(*23*)-thus life as we know it, is truly protein centric. Nucleic acids have limited utility without the framework of proteins around them, controlling, accessing and maintaining them. Lastly, protein constitutes the majority of the cell (75% of a cell is water, the remainder of which is 96% protein, 4% free amino acids and 0.02% nucleic acids, calculated from measurements of muscle cells and estimated to be approximately representative(*24*)).

The following data demonstrate that complex, life forming molecules are not only capable of but are predisposed to self-assembly and self-replication. Unlike the alternative theories on the creation of complex life, these results are reproducible in simple laboratory environments which are representative of real-world environments and conditions present throughout geological time, although more concentrated, without the requirement of complicated coincidental circumstances(*25, 26*).

## Results

Here we demonstrate protein generation from the 20 standard amino acids only, in an aqueous solution, utilizing energy input from the sun (unregulated heat source which fluctuates), or a constant heat source (a culture room at 37°C, for specifics please see the supplementary methods). We present sequence data of the peptides formed; energy input data to determine sufficient energy for amide bond formation, testing for contamination (DNA, RNA and specific conditions including bactericides), quantitative temporal data for >6000 peptides, structural (electron microscopy and 3D modelled) data and machine learning analysis of over 110 characteristic descriptors which enabled us to learn which peptide sequences make good replicators.

Mass spectrometry-based proteomics was the primary mode of analysis. Given that the initial peptide and protein generation was predicted to be stochastic, all spectra were initially identified using De Novo Sequencing (Peaks software(*27*)), with a generated database being used to quantitatively analyse the data using MaxQuant (*28*). Structural characterization was carried out using Scanning Electron Microscopy (SEM) and Transmission Electron Microscopy (TEM). Absence of DNA and RNA was tested using Fluorometric Quantitation (Qubit®, Life Technologies). See full methods in the Supplementary Materials.

### Reproducibility and Breadth of Peptide Synthesis

Mass Spectrometry based proteomic analysis was used for all peptide quantification and sequencing over time. Seven thousand and seventy four peptides were identified with quantitation and the peptide sequence similarity searched using Uniprot Blast(*29*) as MS based database searches do not return partial sequence matches. Of these, 48.3% matched nothing in the described databases (Uniprot TrEmbl), 49.7% have some sequence homology to known proteins and 2% have direct sequence homology to known proteins.

### Machine learning analysis of the peptides generated

Analysis of the peptides present in all samples which displayed increase over the complete time course showed they have an average pI of 4.9, average molecular weight of 1619.7 Da and tend towards helical structures, which are the most common protein structure in nature(*30*). Amino acid utility was measured, and the incorporation of all amino acids was seen, with little deviation in percentage of incorporation compared to that of the Uniprot database (Figure S1).

There are numerous other aspects that describe peptide structure, chemistry, sequence patterns and response to pH and water as a solvent. To effectively describe and interrogate 119 of these characteristics (Supplementary Table S2) we employed machine learning, for the sequence, quantitative and temporal data obtained in this data set.

Briefly, the machine learning focused on 3 outcomes; Peak Intensity: the maximum mass spectrometry signal observed for a given peptide across the full time series, Area Under the Curve (AUC): the integrated abundance across all timepoints, representing cumulative presence in the experiment, and Total Increase: the difference between final and initial intensity values, capturing net growth over the experimental period.

These outcomes, determined as being representative of successful peptide formation and amplification over time, were interrogated against the above mentioned 119 characteristics. These included sequence patterns, asymmetry in charge, spherical tendency, long range and short-range amino acid interactions/proximity, hydrophobicity, dihedral angle composition, and density, among many other measurable attributes.

To allow accurate comparison, we applied SHAP (SHapley Additive exPlanations (*31*)) analysis. This enables identification of which specific molecular properties drive the model’s prediction for any given peptide, providing a principled basis for generating mechanistic hypotheses and prioritising candidates for positive outcomes on quantity of peptide made and longevity. The outcomes of the machine learning analysis are illustrated below (Figure 3).

**Figure 1.**
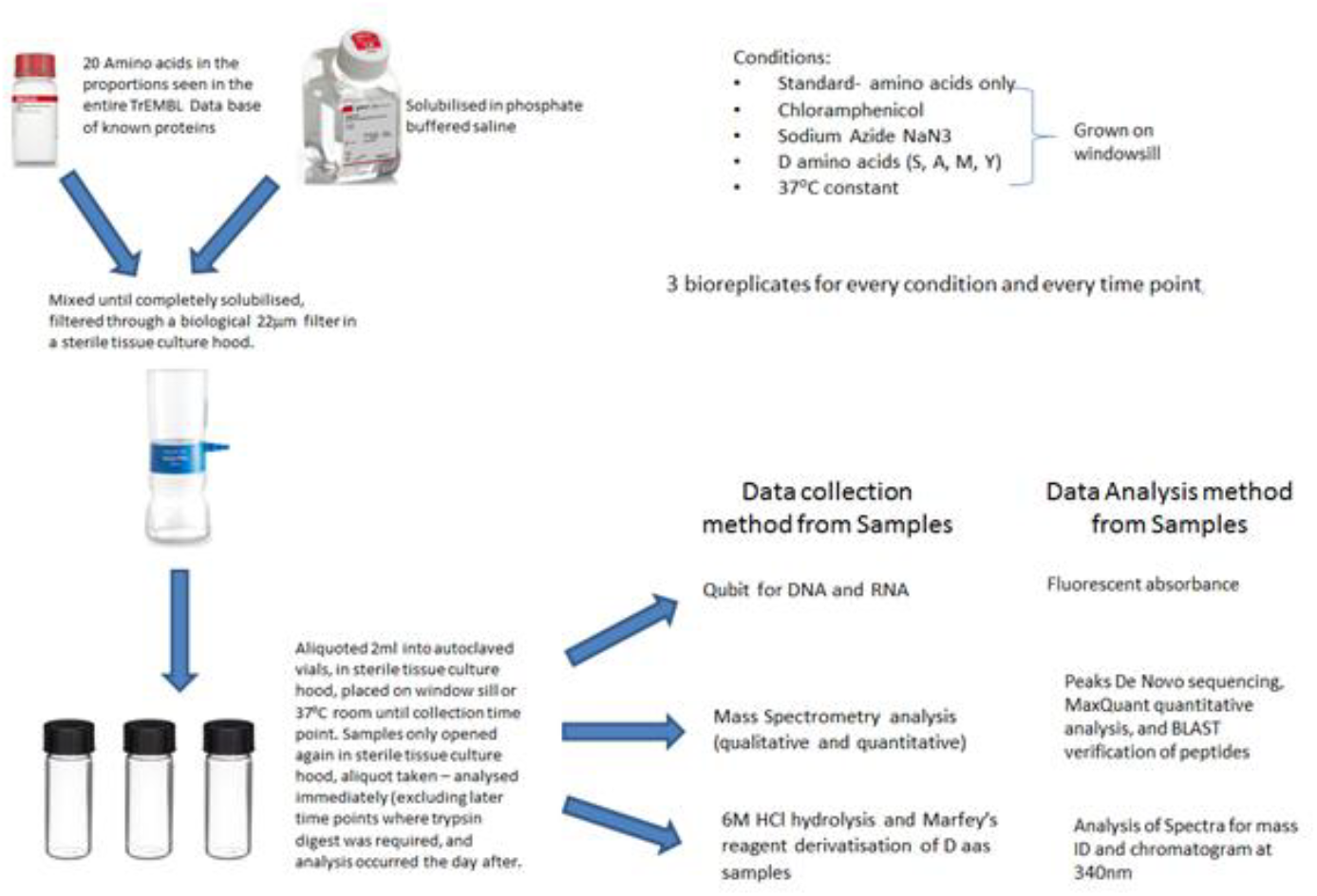
The Experimental Design. The inclusion of additional control samples with sodium azide, chloramphenicol and D amino acids were used to guarantee the absence of biological contamination. Samples were collected at 7-day intervals from T0 to T42. The inclusion of Phosphate Buffered Saline was based on an assumption of required salts and buffering.

**Figure 2.**
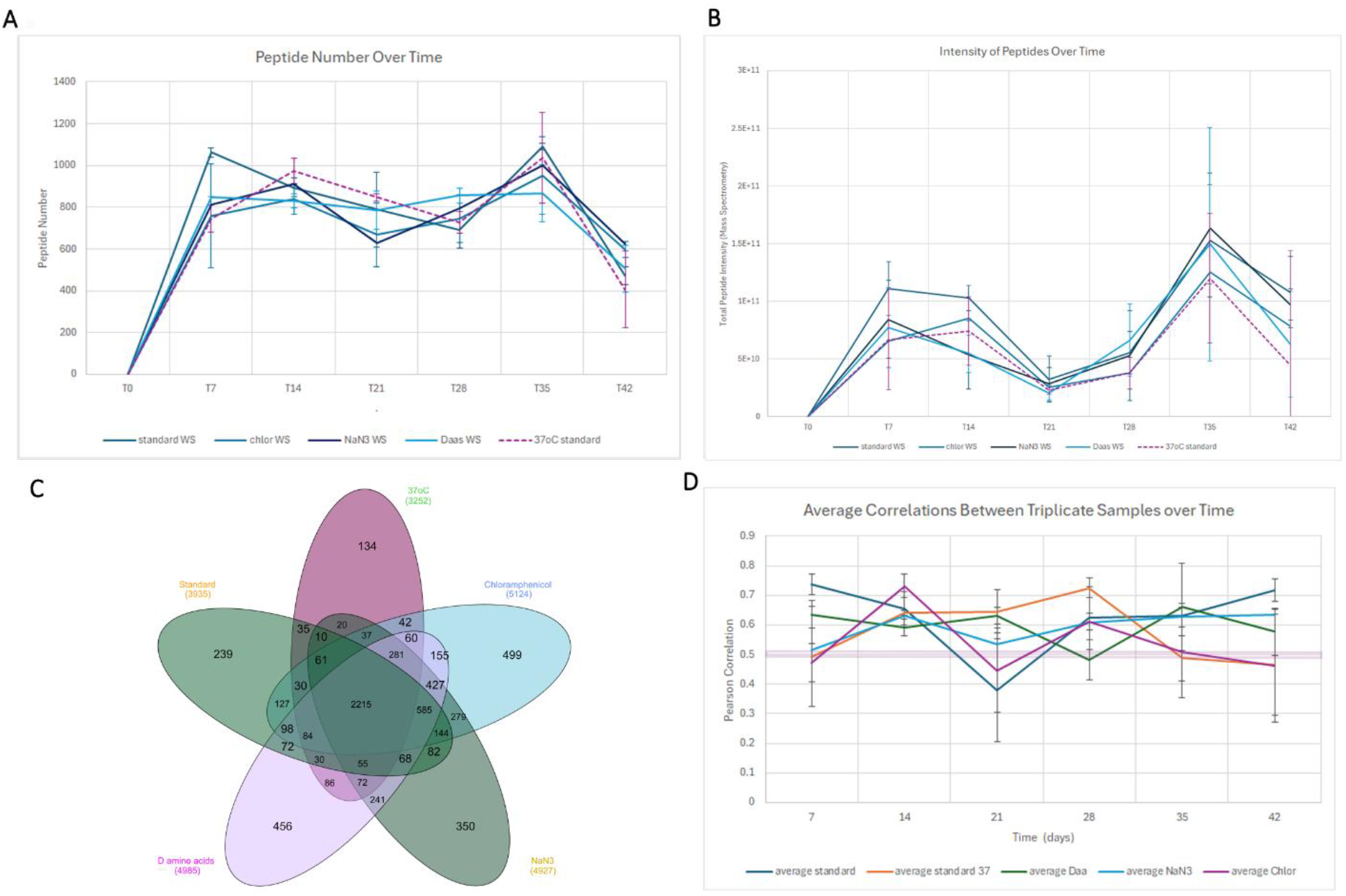
Peptide/protein expression profiles over 42 days in total between all conditions. A. Peptide number seen over time in all conditions, showing between 400 and >1000 different peptides generated over time per condition. B. Peptide intensity derived from mass spectrometry intensity. There is a noted reduction in overall peptide intensity between T14-T28. C. A Venn diagram showing most peptides generated were seen in all conditions, despite each sample being completely independent, as well as showing some unique peptides seen in each condition. The diversity of peptides seen is remarkable with >7000 different peptides seen in samples. D. Pearson correlations over time show the similarity in peptides identified between samples (Pearson correlation value of >0.5) but also quantitatively how the peptides are made in similar amounts as these correlations were calculated from MS intensity data for each peptide and between samples.

**Figure 3.**
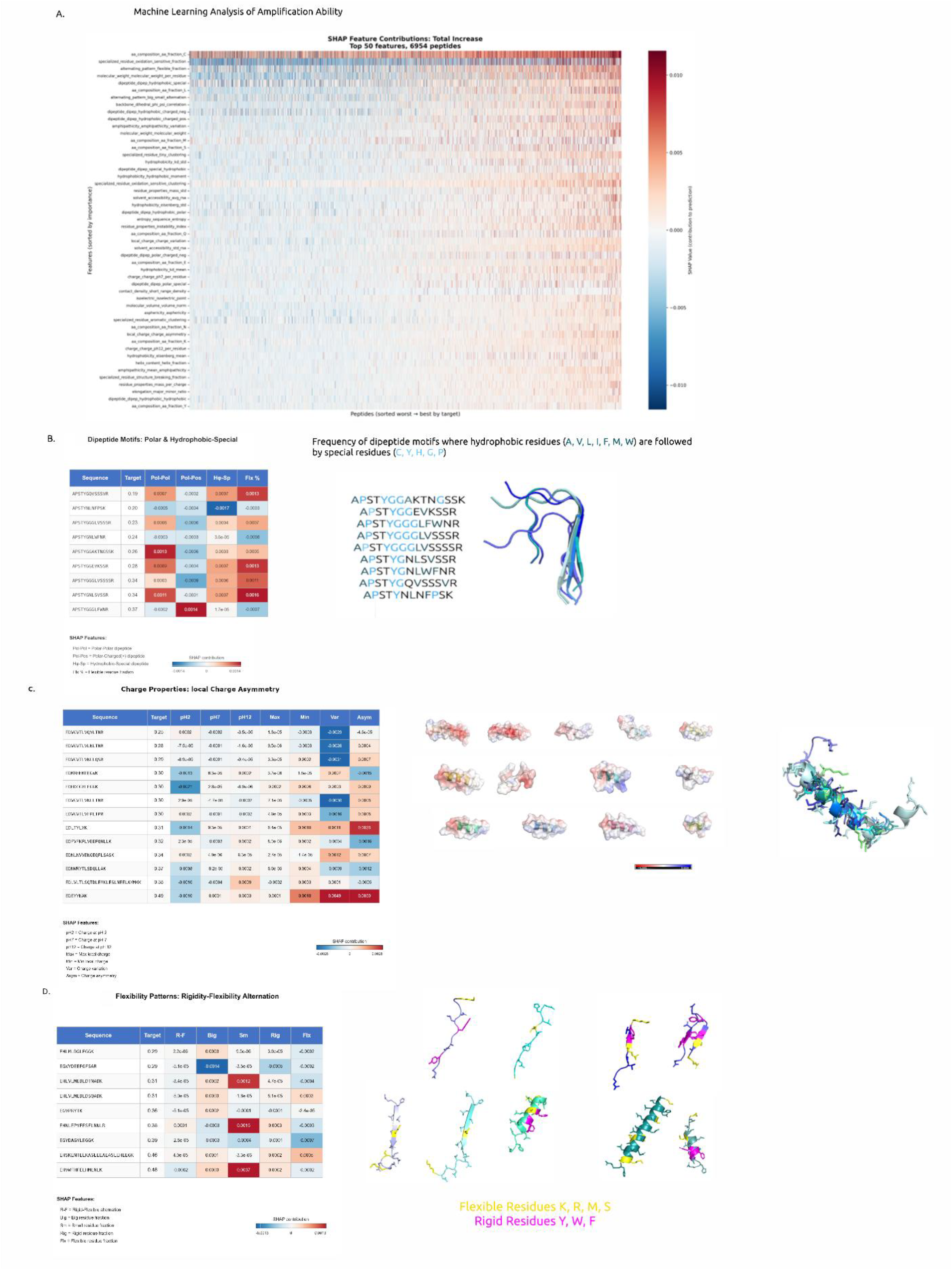
A. A heat map of SHAP values depicting the 119 shape and sequence characteristics impacting the amount and longevity of peptides observed in this experiment (6954 sequences on the X-axis and characteristics on the Y-axis) B. a peptide sequence family (sequence similarity) showing increase in peptide amount related to polar residues being next to hydrophobic residues, and the 3D structures overlaid. C. A Sequence group which showed increase in peptide amount related to local charge asymmetry on the peptide structure, illustrated by the charged surfaces generated in Pymol (*32*). D. A group of structurally very different peptide sequences that show increased peptide amplification related to alternating rigid and flexible residues.

We can identify many aspects which increase the success of peptide formation and amplification, ranging from alternating flexible and rigid residues (Figure 3D) to asymmetry in charge state along the peptide (Figure 3C), to alternating sequences of polar and hydrophobic residues (Figure 3B). The interesting conclusion from these data is that *many* characteristics influence the successful amplification of peptide sequences over time and show this abiogenic templated aqueous peptide amplification is not restricted to only rare sequences (figure 3A). While these data were acquired from trypsin digested proteins, larger proteins were evidenced by the size of proteins and cell-like structures which were later observed via scanning electron microscopy (Figure 5). Despite the required bias to peptides created by trypsinization, we were still able to extract valuable information about peptide formation over time, longevity and suggestions of evolution of a peptide sequence into others (Figure 4B).

**Figure 4.**
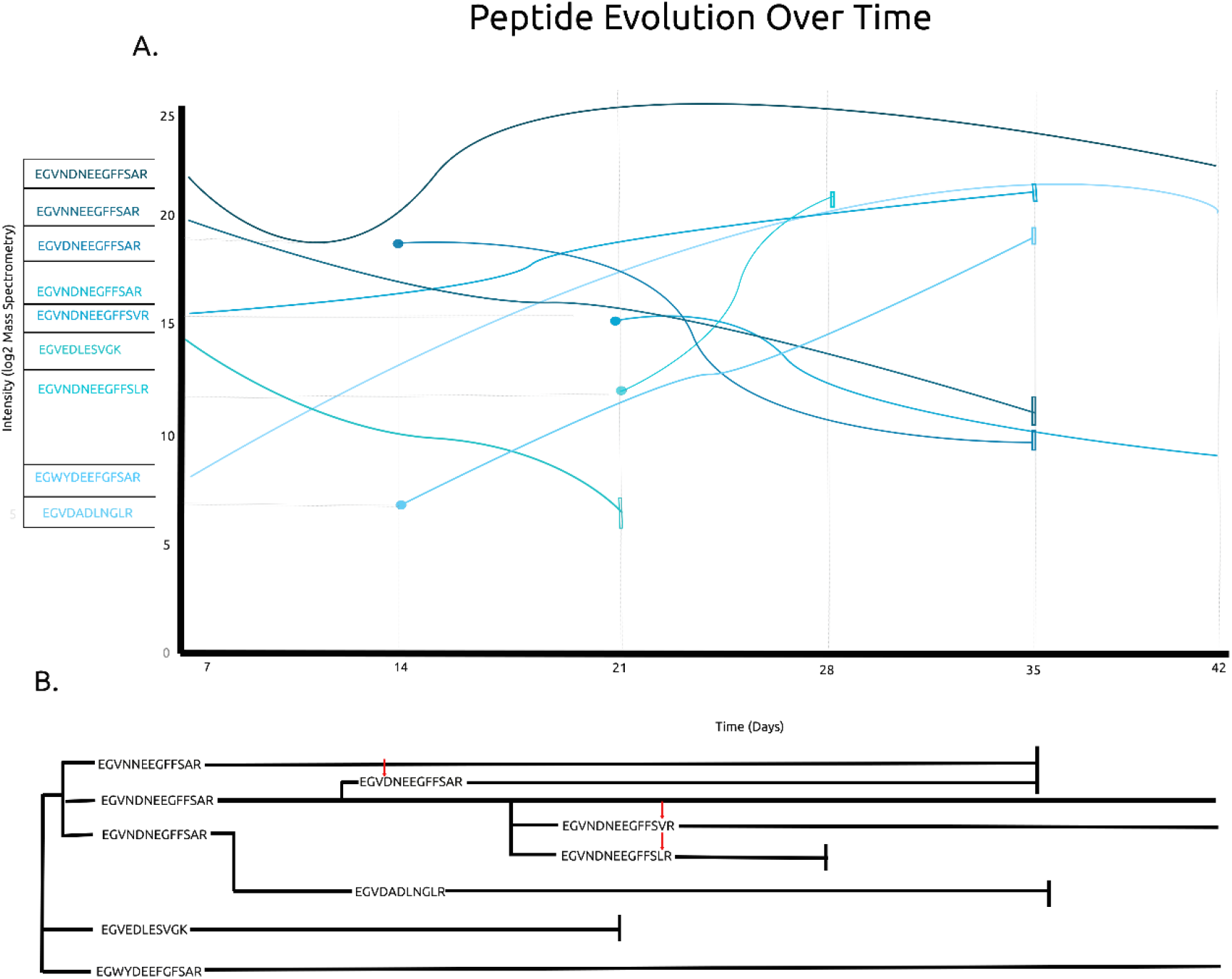
Peptide group evolution. A group of peptides of similar sequence composition can be seen changing through deletion, substitution and full segment miscopy. Unique to this evolution data is the quantification and resolution of time (days and weeks). A. shows the sequence timing (first observed and how long they are observed for) as well as how much of the peptide is there over time, showing decreases and increases (Log2 intensity Mass Spectrometry) B. shows a more traditional evolution tree depicting how a sequence evolved into others and whether the sequences were successful evolutions or whether they ceased to be observed. The sequences line up with the day scale (X axis Figure 4A) shown above, to indicate when the peptide was first observed. Red arrows show the location of the sequence change.

**Figure 5.**
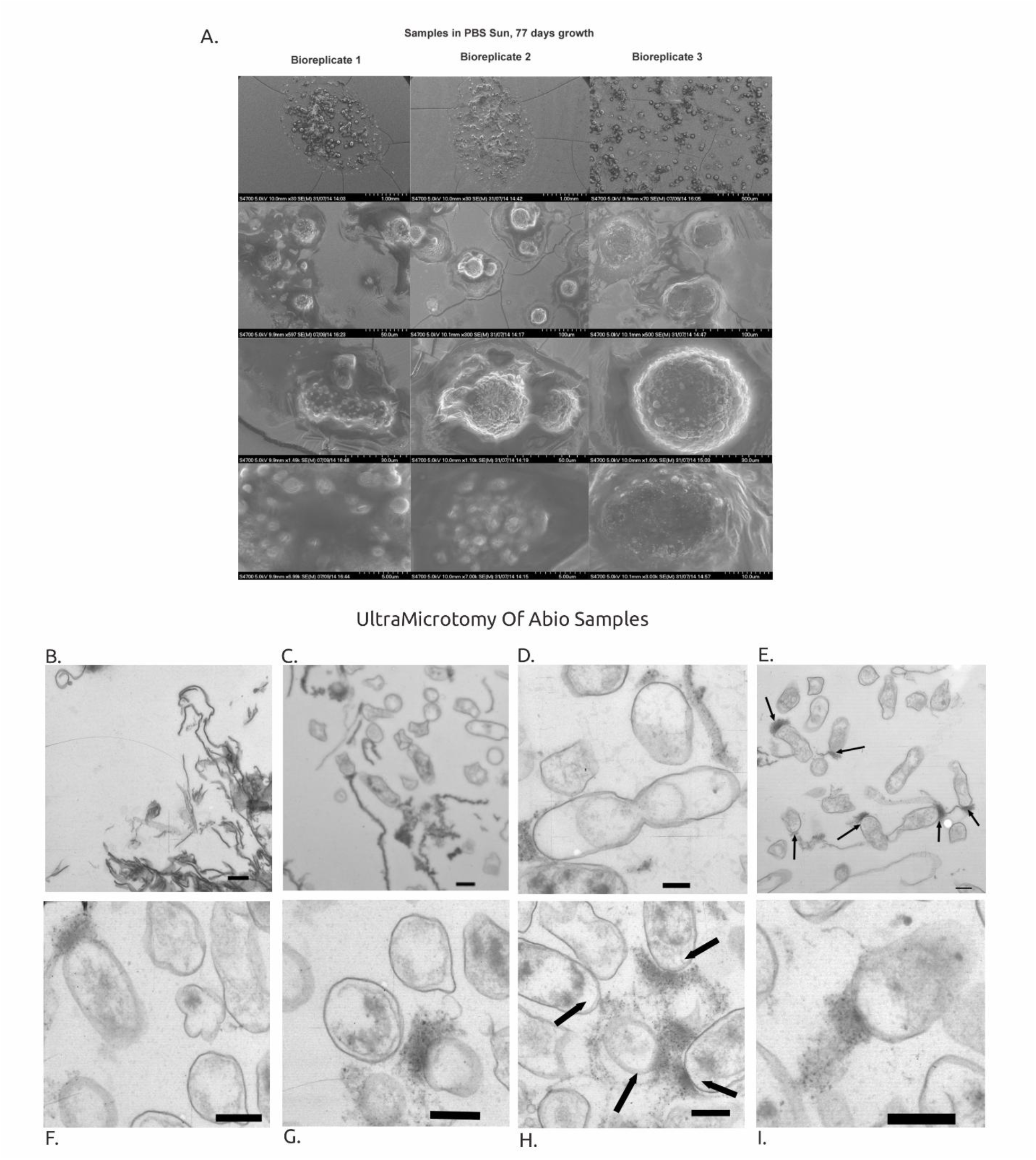
SEM and TEM images of protocells. A.SEM of the larger spherical structures formed at 77 days of abiogenesis. The formation of these structures is reproducible and the variation in sphere size covers the cell size ranges we see today,(*38*) 0.0121mm to 0.1134mm. B. Samples were pelleted by centrifugation, dehydrated in an ethanol series, embedded in resin and ultramicrotomy used to visualize cross sections, including membrane-like structures of the material present in the samples. Protein chains were observed at 60 days of abiogenesis reaction. (bar indicates 437nm) C. Samples from 77 days show presence of cell like cross sections, including a membrane, some completely empty and some with internalized protein. (bar indicates 1346nm) D. Samples from 678 days of abiogenesis. Very clear cell-like structures, with internal, simple, compartmentalization in some cases. The collapse of a ~1536nm cell-like structure into 2 is illustrated. (bar indicates 192nm) E. The aggregation of protein strands at an end on the cell-like structures, seen in ~30% of the structures, marked with arrows. (bar indicates 437nm) F-I. In higher magnification images, there can be seen on the internal side of the membrane a lack of protein staining.

### Peptide Sequence Evolution

The temporal and quantitative nature of the sequence data that was generated, through quantitative proteomics, enabled the ability to observe sequences emerge, mutate and disappear or persist over time. This was possible despite the data being derived from digested proteins, as with proteomics, the peptides are indicative of structure and function (in this case abundance, evolution and longevity) of the protein from which they came. These data suggest certain peptide sequences are preferentially replicating themselves and have structural longevity. Mutations likely arising from miscopies in the replication process, can be observed and the fate of these sequence changes can also be seen Figure 4). The evolution capability observed in these data demonstrates another of Schrodinger’s life characteristics “continual change preceding death”, as well as “the capacity for growth, reproduction”(*33*).

### Imaging – Protein Chains and Pseudo Cells

Samples analysed after 14 days of incubation using HPLC-coupled Mass Spectrometry suggested analytes of a large size. Therefore, samples required breakdown (trypsinisation) of the proteins for mass spectrometry analysis. To understand the larger structures further we employed SEM and TEM to visualize what might be present. Scanning Electron Microscopy revealed spherical structures after relatively short time periods (77 days, Figure 5A), which was followed by ultra-microtomy of 60-, 77-and 678-day samples. This allowed visualization of 70*n*m cross sections of the spherical structures (see supplementary methods).

At the earlier timepoints, long strands of protein can be seen, with chains of protein extending from the long strands. There are also intense spots of protein along the chains that might show new chain formation points, or aggregations (Figure 5B).

Later the circularization of the chains is clear, with the cross sections of the spheres seen in the TEM (Figure5C). The cell-like structures are quite empty when compared to modern cells, with sub-compartments seen. Collapse of larger spheres can be seen (Figure 5D.) which is a known occurrence with increase of size(*34*). The samples analysed after longer periods of Abiogenesis (678 days) exhibit protein aggregation on edges of the cell like structures where the other side of the membrane is devoid of staining, suggesting no protein is present (Figure 5E-I). This could be either an expulsion event or an aggregation event. The encapsulation of life (all life consists of cells) is a primary requirement and the handedness (L or D amino acids and sugars) is attributed to this(*35*). Additionally, encapsulation is thought to enable early metabolism to be maintained, due to differentials across membranes, facilitating flow of nutrients or waste.(*36, 37*)

### Organism/contaminant free reactions

Characterisation of sources of contamination was a priority for these experiments, as aqueous amino acid solutions are ideal environments for microorganism growth. Aside from aseptic sample preparation, and sample analysis, methodologies to detect contamination were employed, including DNA and RNA presence (Figure S1), peptide identification using mass spectrometry, testing of all starting materials (amino acids), and contamination from analytical apparatus. The commercial amino acids used for the experiment were all solubilised in MilliQ water and analysed using MALDI-TOF mass spectrometry. All the samples came back with high purity (supplementary Figures S3A-H), proving they cannot have contributed to the masses and structures we see.

To eliminate possible sources of contamination, all samples were tested for presence of DNA (double stranded) and RNA using the Qubit (Qubit® dsDNA HS Assay Kit, Life Technologies, Q32851, and Qubit® RNA HS Assay Kit, Life Technologies, Q32855). No RNA was detected, while in the samples which changed in colour, low levels of “DNA” were detected. The samples which changed colour were those exposed to UV light, and the colour change was likely due to oxidation of several amino acids. To verify whether this was DNA, the samples and a positive control (*E. coli* lysate) were treated with Benzonase for 60 mins (to degrade any DNA present in the sample). The absorbance seen in the positive control was approximately halved whilst the colour change samples’ absorbance did not change (Supplementary Figure S2) indicating that the proteins in the sample likely auto fluoresce at 530nm (the emission wavelength of Picogreen-the fluorophor used in the Qubit assay).

To identify contamination which may have come from the HPLC column coupled to the mass spectrometer, analytical length blanks were run prior to samples being analysed. The resulting files were searched in the same way as the samples and resulted in 3 Denovo peptides being identified, and 1 peptide match, which was not used in the database for searching and quantitation of the samples.

These tests verify that there was no genetic material present in the samples (which would indicate contamination by a growing organism). Additionally, high numbers of protein hits and full coverage of protein sequences from any given organism were not seen in the proteomics data-which would be the case if there was contamination from an organism in the samples (samples were searched against the entire Uniprot database(*39*)).

Additionally, experimental samples were generated in the presence of Sodium Azide, Chloramphenicol and D amino acids. These additives to the reactions are commonly used bactericides (Sodium Azide and Chloramphenicol) and D amino acids were used as most organisms cannot use these (*40*), or would convert them into L amino acids-which can be tested using Marfey’s reagent (amino acid derivatization to modify the retention time of L vs D amino acids using C18 reversed phase chromatography see supplementary data Methods and Figure S4)(*41*).

In all cases the peptide formation was equivalent (Figure 2C), showing that there was no bacterial contamination, and the reaction is chemistry based, with the additives having no detrimental effect on the peptide formation and amplification outcomes.

### Energy requirements for Amide Bond Formation

To characterize the energy consumption of the reaction required measuring the temperature variations in the natural fluctuating condition (sunlight) and in controlled environments (supplementary Figure S5). Measuring the difference in energy consumption between ambient temperature, aqueous solution only, and aqueous solution with amino acids determined if the conditions were conducive to peptide formation. R. Bruce Martin in 1998(*42*) stated that “in the absence of ionization products, synthesis of peptide bonds would be favoured” further “formation of a di-peptide in an abiogenesis scenario is about 1.2kcal/mol” Martin presented further evidence that this is 8x more difficult than adding an amino acid onto a peptide of any length, and 5x more difficult than joining 2 peptides of at least di-peptide size. To define the amount of energy added to the system, and latterly to compare to control samples (not containing amino acids) we made a bespoke, high accuracy, temperature measurement apparatus (see methods ‘Temperature measurement’). Temperatures were measured every 5 minutes with an accuracy of ±0.0625°C (Figure S5A.) Fluctuations in temperature in the natural sunlight condition were 10.8125°C (2096.7J°C^-1^g^-1^).

To more accurately discern the difference between energy that may be consumed in the dehydration reactions required to make peptide bonds, we used a controlled temperature environment and a control sample of PBS, and the PBS + amino acids solution along with an ambient temperature measurement. We measured every 5 seconds for highly accurate heating and cooling curve characterisation (Figure S5B.) The difference in area under the curve was 28962.97°C^2^s, or 170 °Cs, with the area being less in the PBS amino acids sample (see methods ‘Temperature measurement’ for how the area under the peaks was calculated). The maximum peak in the PBS + amino acids was always less than the control, suggesting energy was always consumed in this system compared to the control. The resulting energy utilised by the PBS + amino acids sample was 1.31kcal, suggesting that in one cycle of heating from 24°C to 34°C and cooling, more than 1mol of di-peptide formation was possible.

## Discussion

From the basic building blocks of proteins, amino acids, in an aqueous solution, free from cellular machinery (RNA, DNA, lipids), not only have proteins been made but structures similar in appearance to cellular membranes and cellular shapes have been observed. These structures agree with Schrodinger’s Paradox, where despite the second law of thermodynamics stating the universe is tending towards disorder; life causes organisation, and therefore a reduction in entropy(*21*).

Through the use of Electron Microscopy (macro structure), fluorescent absorbance (DNA and RNA measurement), novel bespoke computation, physics (system energy consumption), mass spectrometry based proteomics, and Machine Learning assisted data analysis, we were able to image, quantify, and identify peptides not matched to any protein currently described, as well as some homology to known proteins. We were also able to show no bacterial or viral contamination of the samples.

We have evidence of protein/peptide formation and replication (ranging in estimated molecule number from 659 million copies to 187 trillion copies-which statistically cannot happen by chance (1 in 243 quadrillion and 1 in 12 quadrillion respectively to observe a 20mer peptide twice randomly).

The diversity of Denovo sequences identified in such simplistic conditions shows there is potential for the first structural and functional protein products of life. It demonstrates the niche for which life may occur is not a rare or exotic one, rather only amino acids and aqueous conditions, with small temperature inputs. Protein products in an archaeological sense had not been previously widely characterized, but with the advent of Paleo proteomics, this field of analysis has proven proteins to be longer lived than nucleic acids and are currently increasing our knowledge of ancient life(*43-45*).

We suggest this is a method for the genesis of complex life before the system of biology that we understand currently (which relies on cellular machinery to exist and the mechanisms for DNA and RNA to be used as templates for making protein and likely supersede this process due to efficiency) could possibly exist. This process demonstrated vesicle formation, self-replication, and capability for evolution that was selected for through chemical stability. The scenario investigated here was without nucleic acids and lipids which constitute modern cells, but the interaction of these molecules with peptides, showing templated replication has been demonstrated(*46, 47*). Plöger and von Kiedrowski showed that Peptide Nucleic Acids (PNAs) consisting of a peptide backbone and nucleic acids bound to this, enable complementarity and can replicate. This system proved to be self-replicating but was catalysed by the addition of 1-Ethyl-3-(3-dimethylaminopropyl) carbodiimide (EDC), a catalyst to activate carboxyl groups for amide bond formation. Adamala and Szostak showed the interaction of a dipeptide in a lipid vesicle was more easily able to replicate itself in a lipid vesicle than in a free solution, and the presence of the dipeptide further catalysed the incorporation of further lipid molecules, growing the size of the vesicles. Both sets of data highlight the combination of lipids and nucleic acids with peptides enables catalysis, self-replication and increase in size or number of molecules/vesicles.

Mitotic cells not only recover a copy of the genetic material, but also half of a cell membrane, the proteins which package and translate the proteins(*48*) and control mitosis itself. If peptides and proteins have the capability to replicate themselves with fidelity, which we have evidence of here, continuation of information transfer in an enclosed cellular context is possible, assuming there is permeability to amino acids and small peptide fragments in the cellular like structures we have observed. There are the peptide/protein aggregates seen at the terminus of ~30% of the cellular structures observed (Figure 5E-I). The purpose and sequence characterization of these is a future aim, but it does suggest a preference for peptides/proteins to aggregate and preferentially bind in these locations. On the internal side of the membrane, at the site of the peptide/protein aggregations, is a lack of staining, indicating this occurrence could be an expulsion of material, or an attraction to this area due to an imbalance of material (osmosis) on the opposite side of the membrane. These observations are consistent with proposed requirements for membranes/encapsulation to facilitate early metabolism and the handedness (D and L amino acids and sugars) which is consistent in modern life, through the regulation of ions and molecules passing through a membrane, and concentration within a membrane.

Finally, we postulate a theory for what is happening in this abiogenesis of peptides, proteins and cell-like structures. We call this theory the Spontaneous Evolution of Biology (SEB) theory. It incorporates Darwin and Mendel’s theory of natural selection(*49-51*) in a chemical sense, whereby the evolutionary “winners” in this chemical evolution are the protein sequences with longevity derived from function (structural and catalytic self-replicative capability). The initial polymerization of amino acids (Figure 6A) is made possible by the presence of amino acids in solution (0.05g/ml in concentration), aided by sunlight or constant heat, to overcome the required energy to form an amide bond.

**Figure 6.**
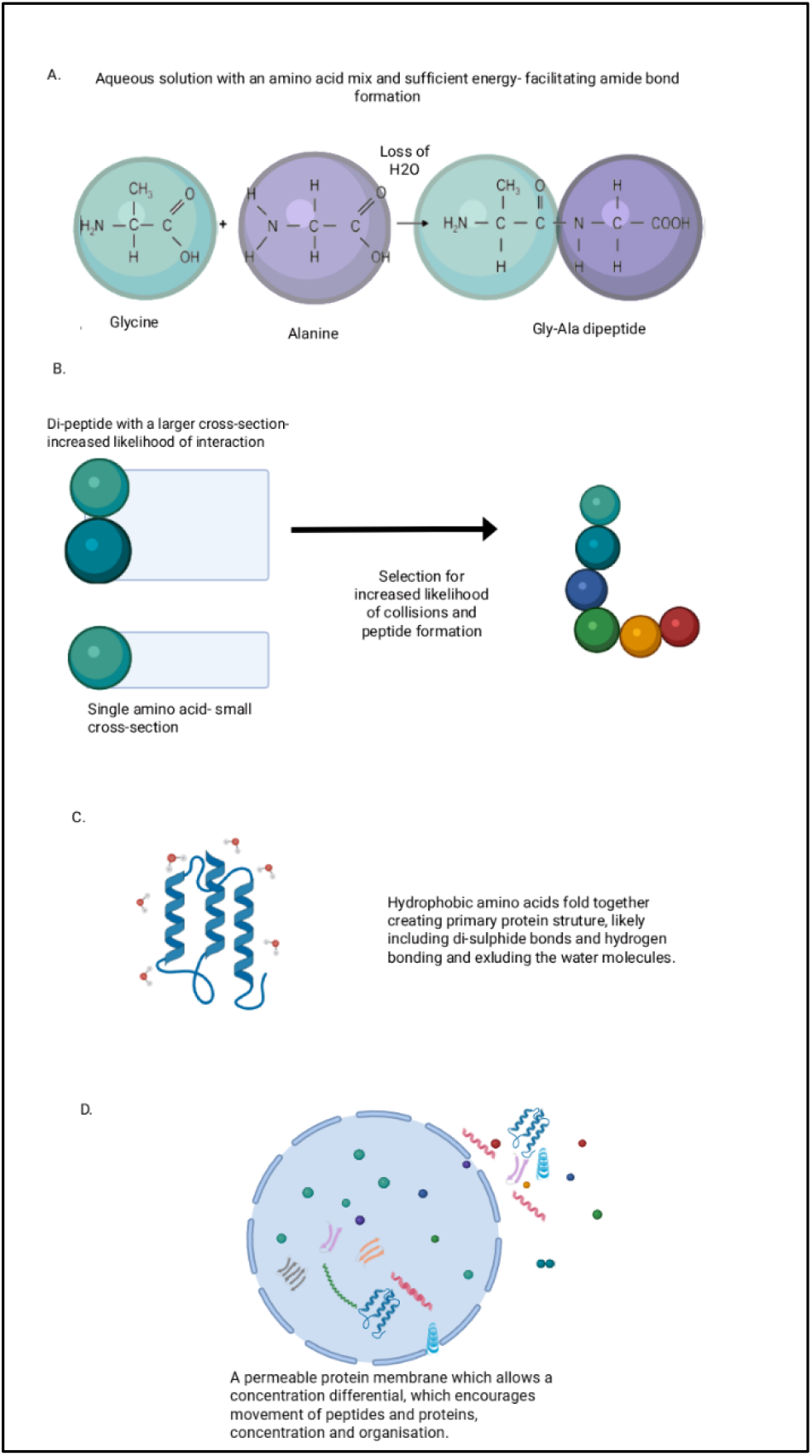
The Spontaneous Evolution of Biology Theory – or SEB theory. A. Amide bond formation in an aqueous solution, provided with adequate energy to facilitate single amino acids to polymerise. B. Further polymerization is selected for due to an increased cross section, increasing the likelihood of interaction with other peptides or single amino acids in the reaction mixture, essentially the peptide works as a catalyst/enzyme. C. More organized structure of longer peptide chains increases longevity and stability-selecting for these structures and enabling catalytic activity (such as self-replication). D. Over longer periods of time cell-like structures were seen, which are selected for as they organize and protect proteins, concentrate amino acids, and enable differential concentrations across the membrane-a driver of energy transfer and generation seen in biology today(*53, 54*).

Subsequent additions of amino acids to existing di-peptides are selected for in 2 ways (Figure 6B). One is an enlarged cross-section increasing the likelihood of collisions with amino acids in solution, facilitating interaction, and secondly the energy required to add amino acids to existing di-peptides is 8x less than the initial amide bond(*42*).

Local chemical environments are a third selection in favour of peptide/protein formation, inducing the folding of the peptide structures, excluding water molecules and forming secondary structures supported with hydrogen bonds and di-sulphide bonds (Figure 6C). This generates structural stability and therefore longevity of peptides/proteins formed.

Finally, the “life” characteristic of reducing entropy by forming spherical cell-like structures (Figure 6D) to organize the proteins occurs. This again protects the structures formed, selecting in favour of these structures. It also presents a physical representation of a structure which resembles the fundamental subunit of complex life-the cell(*52*).

We have empirically shown that peptides and proteins can be spontaneously generated completely independent of cellular machinery and a nucleic acid template. In this scenario, a high amino acid concentration, in an aqueous media with sufficient energy facilitates polymerization of amino acids, replication of peptides with structural longevity, and given time, the formation of cell-like structures.

## Data are available in the supplementary Materials

The authors would like to thank Ass. Prof Andrew Piggott, Department of Molecular Sciences, Macquarie University, for advice on the D and L amino acids using Marfey’s reagent.

## Author contributions

Sara ten Have carried out the experimental procedures and majority of the data analysis. Ewan Macmillan carried out the Machine Learning analysis of the quantitative mass spectrometry peptide data. Rob Kent built the bespoke temperature measurement equipment and carried out the energy use calculations. Alan Prescott carried out the SEM and TEM analysis of the samples. Tim Medway carried out the structural pymol modelling for the figures.

Supplementary Materials:

## Materials and Methods

Tables S1-S2

Figures S1-S4

