## Supplementary data for "The Spontaneous Evolution of Biology"

**Supplementary Materials**

***Sample Preparation***

***Experiment 1***

Commercial amino acids were measured out according to the percentages in table 1 to a total concentration of 1g/100ml. The amino acids were solubilised in either sterile PBS (gibco 1x DPBS 14190-094). Solutions were mixed until complete solubilisation had occurred then passed through 0.1mm stericup filters, and decanted 2ml per autoclaved vial, in a cell culture fume hood with sterilised equipment. Vials were not opened again until an aliquot was required at the specified time point (again all done in a sterile environment). The samples were either analysed that day (if prior to day 14 incubation) or immediately reduced and alkylated, followed by in-solution digestion as described above. For the tryptic digest samples, the samples were reduced and alkylated using DTT and iodoacetamide, followed by tryptic digestion 1:100 ratio of trypsin to protein concentration, for 12 hours at 37^o^C. The digest was halted by the addition of TFA (trifluoroacetic acid) to a final concentration of 0.1%. Samples were desalted over C18 tips, eluted with 80% ACN/0.1% TFA, dried and reconstituted in 40ul 0.1% TFA for injection. An undigested sample aliquot was taken for DNA and RNA analysis and analysed the day of collection also.

Additional samples were left unopened and unanalysed for 60, 77, and 678 days for SEM and TEM analysis.

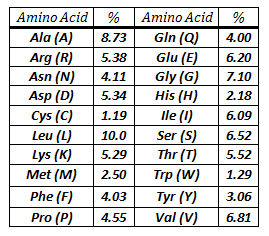

**Table S1**: Total percentage of each of the 20 amino acids commonly found in the UniProtKB/TrEMBL protein database release October 2013 (http://www.ebi.ac.uk/uniprot/TrEMBLstats(*39*)).

**Mass Spectrometry based Proteomic analysis**

Samples were then run on a 120min gradient (2% to 85% acetonitrile) on an EASY-Spray column, 50cm x 75µm ID, PepMap RSLC C18, 2µm, Thermo Scientific (ES803), coupled to a Q-Exactive hybrid Quadrupole Orbitrap mass spectrometer. The full MS survey scan was done to a resolution of 70,000, from 400-1600 m/z. Data dependant acquisition of the top 10 most intense ions was done with an ms/ms resolution of 17,500, and an isolation window of 2.0m/z and a charge exclusion of 1, 8, >8 and unassigned.

The resulting .raw files were processed initially using Peaks 7.0 Denovo sequencing software, and the identified peptides matched to the entire Uniprot database. The denovo peptides which had >50% ALC score, and protein sequences matched from the Uniprot database, were taken to generate a bespoke database for use in MaxQuant 1.5.0.0. The data was searched in MaxQuant with variable modifications Acetyl (N-term), Phospho (STY), Oxidation (M) and Carbomidomethyl (C). It was searched once with Trypsin specific enzyme digestion and with non-specific search (peptide length 5-50aa).

**Scanning Electron Microscopy**

Two microliters of sample were taken (77 Day growth) and applied to silicon wafer pieces and dried at room temperature in a covered petri dish. Specimens were then critically point dried using a BAL-TEC CPD 030. Samples were then mounted on Al stubs using carbon adhesive tabs and coated with 15nm Gold/palladium sputter coating was done using a Cressington 208HR sputter coater. Specimens were examined using a Hitachi S-4700 operating at an accelerating voltage of 5kV

**Transmission Electron Microscopy**

**Direct TEM**

Initially samples were analysed without processing of any sort. Five microliters of sample were applied to the grid, stained with 0.05M Uranyl acetate in 0.05M HCl. Specimens were examined using the JEOL JEM1200 EX

**Ultra-Microtomy TEM**

To elucidate (if possible) the internal structure of the spherical structures observed we used Ultra-microtomy SEM. Ten millilitres of sample were spun down at room temperature at 1000rpm for 1 hour. The supernatant was removed from the pellet and the pellet re-suspended in 500μl 1.4% w/v ultra-pure agarose (Invitrogen 16500-500). The resulting solidified agarose was cubed in 1mm^3^ and treated with 1% osmium tetroxide for 1 hour at room temperature. Samples were then washed with distilled water and treated with 1% uranyl acetate overnight at room temperature. The samples were then gradually dehydrated with gradually increasing concentrations of methanol and finally propylene oxide. The dehydrated samples were embedded in epoxy resin and sectioned. Specimens were examined using the JEOL JEM1200 EX

**Temperature measurement**

Bespoke instrumentation was created for the purpose of measuring temperature precisely and to record it at specified time intervals over a long period of time. This was achieved by coupling Maxim IC DS18B20+ temperature sensors (product number, accuracy of ±0.0625^o^C) to Arduino Leonardo clone single board micro controllers (<http://arduino.cc/> Code: A000057), with a real time clock, Maxim DS1307. The output was then read to initially SD cards then directly via USB to a bespoke python application with live plotting interface. Full details of the project including the functioning code can be accessed here (<http://www.instructables.com/id/Quick-Easy-Temperature-Loggers/?fb_action_ids=10204855266357922&fb_action_types=og.shares> ). For the controlled heating experiment measurements were taken every 5 seconds, resulting in over 145,000 data points.

Calculation of the energy required to make amide bonds

The area under the curve was calculates using the Trapezoidal Uniform Grid method
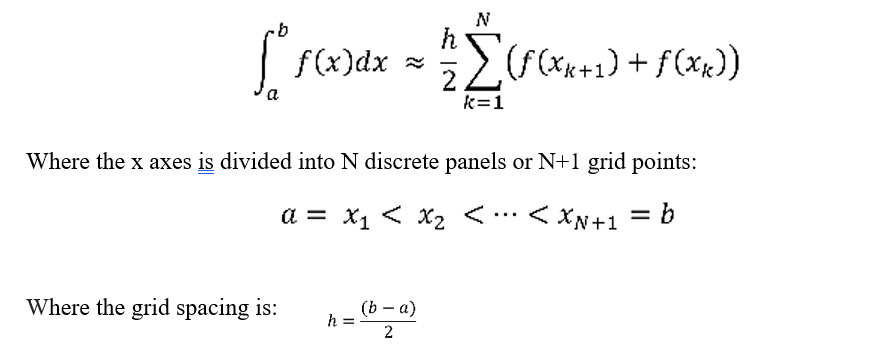

**DNA and RNA Detection**

The Qubit commercial reagents were used according to the instructions for Qubit® dsDNA HS Assay Kit, Life Technologies, Q32851, and Qubit® RNA HS Assay Kit, Life Technologies, Q32855. Twenty microliters of sample were used for the assays.

The samples were measured on the Qubit® 2.0 Fluorometer, Q32866.

**Marfey’s Analysis of D Amino Acids Composition**

One hundred microliters of each bio replicate of the D amino acids samples (T28) and the standard samples (T28) were pooled together resulting in 1 sample for each condition. The samples were rotary evaporated to dryness. One mililtre of 6M HCl was then added to the samples, and they were heated at 155^o^C for 80 minutes. The samples were then rotary evaporated to dryness. Samples were re-constituted with 300μl of MilliQ water, and pH adjusted (to >5) with 3μl of 10M NaOH. One hundred microliters were combined with 200μl of Marfey’s reagent(*41*) (10mg/ml in acetone) and 40μl of 1M ammonium bicarbonate and gently shaken at 40^o^C for 1 hour. The reaction was quenched with 20μl of 2M HCl. Five microliters of sample were injected onto a 50cm pepmap C18 column, and the UV absorbance measured at 340nm. Mass Acquisition was performed using a Thermo Orbitrp XL, top 5 ions selected for ms/ms with an ms resolution of 60,000 (FTMS), scanning from 200m/z- 1800m/z.

**Machine Learning Analysis**

**Target Metrics**

Three prevalence metrics, each capturing a different aspect of peptide behaviour in the abiogenesis experiments, were used as proxies for templating potential:

Peak Intensity: the maximum mass spectrometry signal observed for a given peptide across the full time series.

Area Under the Curve (AUC): the integrated abundance across all timepoints, representing cumulative presence in the experiment.

Total Increase: the difference between final and initial intensity values, capturing net growth over the experimental period.

Raw metric values were first averaged across experimental replicates, then log-transformed to normalise the strongly right-skewed distributions typical of mass spectrometry data. Finally, values were standardised to z-scores (mean = 0, standard deviation = 1) to place all three metrics on a comparable scale. For the classification task, peptides scoring above the 75th percentile on each metric were labelled as positive cases (the top quartile), with inverse-frequency class weighting applied during model training to account for the resulting 3:1 class imbalance.

**Feature Engineering**

A total of 133 molecular descriptors were computed for each peptide sequence. These fell into two broad categories.

**Sequence-Based Features**

These features were derived directly from the amino acid sequence and required no structural prediction. They included standard physicochemical properties such as molecular weight, isoelectric point, net charge at pH 7, GRAVY hydrophobicity score, instability index, and aromaticity. Amino acid composition was captured as the frequency of each of the 20 standard amino acids, supplemented by dipeptide frequencies (counts of adjacent amino acid pairs, e.g. Lys-Arg, Pro-Pro). Position-specific features describing the physicochemical properties of the N-terminal and C-terminal residues were also included.

**Structure-Based Features**

Structural descriptors were obtained from three-dimensional models predicted using AlphaFold2 via ColabFold. These included secondary structure content (helix, sheet, and coil fractions), solvent accessibility (proportions of exposed versus buried residues), compactness metrics (radius of gyration, contact density), and the pLDDT confidence score, which reflects the predicted reliability of local structural predictions on a per-residue basis.

**Feature Preprocessing**

Prior to model training, the feature set was refined through several preprocessing steps. Features with near-zero variance (threshold < 0.01) were removed, as these carry negligible discriminative information. Highly correlated feature pairs (Pearson r > 0.98) were identified, and one member of each pair was iteratively dropped to reduce redundancy. Any remaining missing values were imputed using the median of the respective feature. All features were then z-score standardised using statistics computed exclusively from the training set, to prevent information leakage from the validation and test partitions.

After preprocessing, the final feature set comprised 119 features.

**Model Development**

The dataset was divided into training (68%), validation (12%), and held-out test (20%) partitions, with stratified sampling applied for the classification task to maintain consistent class proportions across splits.

We employed XGBoost, a gradient-boosted decision tree algorithm widely used in tabular prediction tasks. In brief, gradient boosting works by training an ensemble of decision trees sequentially, where each successive tree is fitted to the residual errors of the preceding ensemble, thereby progressively refining the predictions. Early stopping on the validation set was used to halt training once performance ceased to improve, guarding against overfitting.

Hyperparameters were optimised via random search, evaluating 500 configurations for the regression models and 500 for the classification models. The search space covered tree depth (3–10), learning rate (0.01–0.1), L1 and L2 regularisation strength, and subsampling rates. Regression models were selected on the basis of validation set Pearson correlation, while classification models were selected by ROC-AUC (the area under the receiver operating characteristic curve, which measures the model’s ability to rank positive cases above negative ones).

**SHAP Analysis**

To move beyond aggregate feature importance and understand individual predictions, we applied SHAP (SHapley Additive exPlanations) analysis(*31*). SHAP is rooted in cooperative game theory: it assigns each feature a contribution score for every prediction by computing the marginal effect of including or excluding that feature across all possible feature combinations. This decomposition enables identification of which specific molecular properties drive the model’s prediction for any given peptide, providing a principled basis for generating mechanistic hypotheses and prioritising candidates for experimental follow-up.

**Table S2**- Machine Learning Characteristics used to interrogate amplification characteristics

| **Feature Name** | **Definition** |
| --- | --- |
| chain_length_chain_length | Number of amino acid residues in the peptide sequence (sequence length) |
| chain_length_chain_length_log | Log-transformed sequence length; reduces the influence of very long sequences |
| molecular_weight_molecular_weight | Total molecular mass of the peptide in Daltons (Da), calculated from amino acid composition |
| molecular_weight_molecular_weight_per_residue | Average molecular mass per amino acid residue (total MW / sequence length) |
| aa_composition_aa_fraction_A | Fraction of Alanine (A) residues in the sequence (0-1 scale) |
| aa_composition_aa_fraction_C | Fraction of Cysteine (C) residues in the sequence (0-1 scale) |
| aa_composition_aa_fraction_D | Fraction of Aspartic acid (D) residues in the sequence (0-1 scale) |
| aa_composition_aa_fraction_E | Fraction of Glutamic acid (E) residues in the sequence (0-1 scale) |
| aa_composition_aa_fraction_F | Fraction of Phenylalanine (F) residues in the sequence (0-1 scale) |
| aa_composition_aa_fraction_G | Fraction of Glycine (G) residues in the sequence (0-1 scale) |
| aa_composition_aa_fraction_H | Fraction of Histidine (H) residues in the sequence (0-1 scale) |
| aa_composition_aa_fraction_I | Fraction of Isoleucine (I) residues in the sequence (0-1 scale) |
| aa_composition_aa_fraction_K | Fraction of Lysine (K) residues in the sequence (0-1 scale) |
| aa_composition_aa_fraction_L | Fraction of Leucine (L) residues in the sequence (0-1 scale) |
| aa_composition_aa_fraction_M | Fraction of Methionine (M) residues in the sequence (0-1 scale) |
| aa_composition_aa_fraction_N | Fraction of Asparagine (N) residues in the sequence (0-1 scale) |
| aa_composition_aa_fraction_P | Fraction of Proline (P) residues in the sequence (0-1 scale) |
| aa_composition_aa_fraction_Q | Fraction of Glutamine (Q) residues in the sequence (0-1 scale) |
| aa_composition_aa_fraction_R | Fraction of Arginine (R) residues in the sequence (0-1 scale) |
| aa_composition_aa_fraction_S | Fraction of Serine (S) residues in the sequence (0-1 scale) |
| aa_composition_aa_fraction_T | Fraction of Threonine (T) residues in the sequence (0-1 scale) |
| aa_composition_aa_fraction_V | Fraction of Valine (V) residues in the sequence (0-1 scale) |
| aa_composition_aa_fraction_W | Fraction of Tryptophan (W) residues in the sequence (0-1 scale) |
| aa_composition_aa_fraction_Y | Fraction of Tyrosine (Y) residues in the sequence (0-1 scale) |
| dipeptide_dipep_hydrophobic_hydrophobic | Frequency of dipeptide motifs where hydrophobic residues (A, V, L, I, F, M, W) are followed by hydrophobic residues (A, V, L, I, F, M, W) |
| dipeptide_dipep_hydrophobic_polar | Frequency of dipeptide motifs where hydrophobic residues (A, V, L, I, F, M, W) are followed by polar uncharged residues (S, T, N, Q) |
| dipeptide_dipep_polar_charged_neg | Frequency of dipeptide motifs where polar uncharged residues (S, T, N, Q) are followed by negatively charged residues (D, E) |
| dipeptide_dipep_charged_neg_polar | Frequency of dipeptide motifs where negatively charged residues (D, E) are followed by polar uncharged residues (S, T, N, Q) |
| dipeptide_dipep_polar_hydrophobic | Frequency of dipeptide motifs where polar uncharged residues (S, T, N, Q) are followed by hydrophobic residues (A, V, L, I, F, M, W) |
| dipeptide_dipep_hydrophobic_charged_neg | Frequency of dipeptide motifs where hydrophobic residues (A, V, L, I, F, M, W) are followed by negatively charged residues (D, E) |
| dipeptide_dipep_polar_polar | Frequency of dipeptide motifs where polar uncharged residues (S, T, N, Q) are followed by polar uncharged residues (S, T, N, Q) |
| dipeptide_dipep_polar_charged_pos | Frequency of dipeptide motifs where polar uncharged residues (S, T, N, Q) are followed by positively charged residues (K, R) |
| dipeptide_dipep_hydrophobic_special | Frequency of dipeptide motifs where hydrophobic residues (A, V, L, I, F, M, W) are followed by special residues (C, Y, H, G, P) |
| dipeptide_dipep_special_special | Frequency of dipeptide motifs where special residues (C, Y, H, G, P) are followed by special residues (C, Y, H, G, P) |
| dipeptide_dipep_special_hydrophobic | Frequency of dipeptide motifs where special residues (C, Y, H, G, P) are followed by hydrophobic residues (A, V, L, I, F, M, W) |
| dipeptide_dipep_charged_neg_hydrophobic | Frequency of dipeptide motifs where negatively charged residues (D, E) are followed by hydrophobic residues (A, V, L, I, F, M, W) |
| dipeptide_dipep_special_charged_neg | Frequency of dipeptide motifs where special residues (C, Y, H, G, P) are followed by negatively charged residues (D, E) |
| dipeptide_dipep_hydrophobic_charged_pos | Frequency of dipeptide motifs where hydrophobic residues (A, V, L, I, F, M, W) are followed by positively charged residues (K, R) |
| dipeptide_dipep_charged_neg_special | Frequency of dipeptide motifs where negatively charged residues (D, E) are followed by special residues (C, Y, H, G, P) |
| dipeptide_dipep_special_charged_pos | Frequency of dipeptide motifs where special residues (C, Y, H, G, P) are followed by positively charged residues (K, R) |
| dipeptide_dipep_charged_pos_charged_neg | Frequency of dipeptide motifs where positively charged residues (K, R) are followed by negatively charged residues (D, E) |
| dipeptide_dipep_charged_neg_charged_neg | Frequency of dipeptide motifs where negatively charged residues (D, E) are followed by negatively charged residues (D, E) |
| dipeptide_dipep_special_polar | Frequency of dipeptide motifs where special residues (C, Y, H, G, P) are followed by polar uncharged residues (S, T, N, Q) |
| dipeptide_dipep_polar_special | Frequency of dipeptide motifs where polar uncharged residues (S, T, N, Q) are followed by special residues (C, Y, H, G, P) |
| dipeptide_dipep_charged_pos_special | Frequency of dipeptide motifs where positively charged residues (K, R) are followed by special residues (C, Y, H, G, P) |
| dipeptide_dipep_charged_neg_charged_pos | Frequency of dipeptide motifs where negatively charged residues (D, E) are followed by positively charged residues (K, R) |
| dipeptide_dipep_charged_pos_polar | Frequency of dipeptide motifs where positively charged residues (K, R) are followed by polar uncharged residues (S, T, N, Q) |
| dipeptide_dipep_charged_pos_hydrophobic | Frequency of dipeptide motifs where positively charged residues (K, R) are followed by hydrophobic residues (A, V, L, I, F, M, W) |
| dipeptide_dipep_charged_pos_charged_pos | Frequency of dipeptide motifs where positively charged residues (K, R) are followed by positively charged residues (K, R) |
| entropy_sequence_entropy | Shannon entropy of amino acid composition; measures sequence diversity (0 = single amino acid type, ~4.3 bits = uniform distribution of all 20 amino acids) |
| isoelectric_isoelectric_point | Isoelectric point (pI): the pH at which the peptide has zero net electrical charge (range 0-14) |
| charge_charge_ph2_per_residue | Net electrical charge per residue at pH 2 (acidic conditions); positive values indicate cationic peptide |
| charge_charge_ph7_per_residue | Net electrical charge per residue at pH 7 (physiological pH); indicates peptide polarity at neutral pH |
| charge_charge_ph12_per_residue | Net electrical charge per residue at pH 12 (basic conditions); negative values indicate anionic peptide |
| local_charge_max_local_charge | Maximum local charge within a sliding window along the sequence; indicates regions of high charge density |
| local_charge_min_local_charge | Minimum local charge within a sliding window; indicates regions of negative charge accumulation |
| local_charge_charge_variation | Standard deviation of local charges along the sequence; high values indicate uneven charge distribution |
| local_charge_charge_asymmetry | Difference in charge between N-terminal and C-terminal regions; measures charge polarity along the peptide |
| hydrophobicity_kd_mean | Mean hydrophobicity using the Kyte-Doolittle scale; positive values indicate hydrophobic character |
| hydrophobicity_kd_std | Standard deviation of Kyte-Doolittle hydrophobicity along the sequence; measures hydrophobicity variation |
| hydrophobicity_eisenberg_mean | Mean hydrophobicity using the Eisenberg consensus scale; alternative measure of overall hydrophobic character |
| hydrophobicity_eisenberg_std | Standard deviation of Eisenberg hydrophobicity; measures the variation in hydrophobicity along the sequence |
| hydrophobicity_hydrophobic_moment | Hydrophobic moment: measures amphipathicity (segregation of hydrophobic/hydrophilic faces), important for membrane interaction potential |
| amphipathicity_max_amphipathicity | Maximum local amphipathicity score; indicates peak segregation of hydrophobic and hydrophilic residues |
| amphipathicity_mean_amphipathicity | Average amphipathicity along the sequence; overall measure of hydrophobic/hydrophilic face separation |
| amphipathicity_amphipathicity_variation | Standard deviation of local amphipathicity; measures consistency of amphipathic character |
| periodic_patterns_repeats_2_density | Density of 2-residue repeating patterns (e.g., XYXY); indicates sequence periodicity |
| periodic_patterns_repeats_3_density | Density of 3-residue repeating patterns (e.g., XYZXYZ); common in structural motifs |
| periodic_patterns_repeats_4_density | Density of 4-residue repeating patterns; may indicate regular secondary structure |
| periodic_patterns_repeat_ratio_2_3 | Ratio of 2-mer to 3-mer repeat densities; characterizes the dominant repeat pattern type |
| residue_properties_average_mass | Average molecular mass of individual amino acid residues in Daltons |
| residue_properties_mass_std | Standard deviation of residue masses; indicates diversity in residue sizes |
| residue_properties_instability_index | Instability index predicting in vivo half-life; values >40 suggest unstable proteins |
| residue_properties_mass_per_charge | Ratio of total mass to number of charged residues (log-transformed); indicates charge density |
| positional_n_term_charged_frac | Fraction of charged residues (D, E, K, R) in the N-terminal region (first 5 residues) |
| positional_c_term_charged_frac | Fraction of charged residues (D, E, K, R) in the C-terminal region (last 5 residues) |
| positional_n_term_hydrophobic_frac | Fraction of hydrophobic residues in the N-terminal region; affects membrane insertion |
| positional_c_term_hydrophobic_frac | Fraction of hydrophobic residues in the C-terminal region |
| positional_terminal_charge_bias | Asymmetry in charge distribution between N-terminal and C-terminal regions |
| specialized_residue_aromatic_fraction | Fraction of aromatic residues (Phe, Trp, Tyr); important for π-stacking and hydrophobic interactions |
| specialized_residue_aromatic_clustering | Clustering coefficient of aromatic residues; measures tendency to cluster together in the sequence |
| specialized_residue_aromatic_max_density | Maximum local density of aromatic residues in a sliding window |
| specialized_residue_oxidation_sensitive_fraction | Fraction of oxidation-sensitive residues (Cys, Met, Trp); indicates susceptibility to oxidative damage |
| specialized_residue_oxidation_sensitive_clustering | Clustering of oxidation-sensitive residues; clustered residues may be more vulnerable to oxidation |
| specialized_residue_oxidation_sensitive_max_density | Maximum local density of oxidation-sensitive residues |
| specialized_residue_structure_breaking_fraction | Fraction of structure-breaking residues (Gly, Pro); these disrupt regular secondary structures |
| specialized_residue_structure_breaking_clustering | Clustering of Gly and Pro residues; affects flexibility and structural disorder |
| specialized_residue_structure_breaking_max_density | Maximum local density of structure-breaking residues |
| specialized_residue_tiny_fraction | Fraction of tiny residues (Ala, Gly, Ser); small side chains allow tight packing |
| specialized_residue_tiny_clustering | Clustering coefficient of tiny residues in the sequence |
| specialized_residue_tiny_max_density | Maximum local density of tiny residues |
| alternating_pattern_big_small_alternation | Score measuring alternation between big (F, W, Y, K, R, M) and small (A, G, C, S) residues |
| alternating_pattern_rigid_flexible_alternation | Score measuring alternation between rigid (W, Y, F) and flexible (R, K, M, S) residues |
| alternating_pattern_big_fraction | Fraction of bulky residues (Phe, Trp, Tyr, Lys, Arg, Met) in the sequence |
| alternating_pattern_small_fraction | Fraction of small residues (Ala, Gly, Cys, Ser) in the sequence |
| alternating_pattern_rigid_fraction | Fraction of conformationally rigid residues (Trp, Tyr, Phe) |
| alternating_pattern_flexible_fraction | Fraction of conformationally flexible residues (Arg, Lys, Met, Ser) |
| helix_content_helix_fraction | Fraction of residues in α-helical conformation (predicted from 3D structure using DSSP) |
| sheet_content_sheet_fraction | Fraction of residues in β-sheet conformation (extended strand structures) |
| turn_content_turn_fraction | Fraction of residues in turn conformations (reverse direction of the peptide chain) |
| coil_content_coil_fraction | Fraction of residues in random coil (unstructured/flexible regions) |
| solvent_accessibility_avg_rsa | Average relative solvent accessibility (0-1); measures how exposed residues are to solvent |
| solvent_accessibility_std_rsa | Standard deviation of solvent accessibility; indicates variation in exposure along the sequence |
| residue_exposure_buried_fraction | Fraction of residues that are buried (RSA < 0.2); indicates core/interior residues |
| residue_exposure_exposed_fraction | Fraction of residues that are solvent-exposed (RSA > 0.5); surface-accessible residues |
| contact_order_0 | Relative contact order: average sequence separation of contacting residues, normalized by sequence length; higher values indicate more complex folding topology |
| long_range_contact_lr_contact_density | Density of long-range contacts (>5 residues apart); indicates tertiary structure complexity |
| long_range_contact_lr_contact_distribution | Coefficient of variation in long-range contact distribution; measures contact uniformity |
| interaction_density_local_density | Density of local contacts (<4 residues apart in sequence); indicates local structural compactness |
| interaction_density_nonlocal_density | Density of non-local contacts (≥4 residues apart); indicates long-range structural interactions |
| contact_density_short_range_density | Density of short-range atomic contacts (<5 residues apart in sequence) |
| contact_density_long_range_density | Density of long-range atomic contacts (≥5 residues apart); key indicator of tertiary structure |
| radius_gyration_rg_norm | Normalized radius of gyration: RMS distance of atoms from center of mass, indicates overall compactness (normalized by 30 Å) |
| asphericity_asphericity | Asphericity parameter (0-1): deviation from spherical shape; 0 = perfect sphere, 1 = maximally elongated |
| elongation_major_minor_ratio | Ratio of major to minor principal axes; values >1 indicate elongated shape |
| elongation_planarity | Planarity: ratio of intermediate to major axis; indicates whether structure is planar or globular |
| molecular_volume_volume_norm | Molecular volume (convex hull) normalized by 1000 Å³; overall 3D space occupied by the peptide |
| backbone_dihedral_phi_psi_correlation | Correlation between φ and ψ backbone torsion angles; indicates regularity of backbone conformation |
| backbone_dihedral_ramachandran_outliers | Fraction of residues with φ/ψ angles outside allowed Ramachandran regions (transformed); indicates structural strain or unusual conformations |

Amino Acid Utility

|  | Abio % | Uniprot 2026 |
| --- | --- | --- |
| A | 6.68318% | 8.69% |
| C | 1.1984% | 1.42% |
| D | 6.16703% | 5.44% |
| E | 7.20526% | 6.28% |
| F | 4.01495% | 3.84% |
| G | 6.16259% | 7.07% |
| H | 2.27964% | 2.3% |
| I | 0.034113% | 5.33% |
| K | 7.14296% | 5.05% |
| L | 9.67474% | 9.76% |
| M | 3.217% | 2.32% |
| N | 6.22191% | 3.87% |
| P | 4.35015% | 5.15% |
| Q | 5.1036% | 3.93% |
| R | 5.18962% | 5.81% |
| S | 7.08512% | 7.2% |
| T | 5.59601% | 5.6% |
| V | 6.23526% | 6.75% |
| W | 2.06754% | 1.3% |
| Y | 4.37091% | 2.85% |

Figure S1. Utility of amino acids compared to the Uniprot January 2026 database. The major discrepancy between Isoleucine seen in this data set is due to mass spectrometry not being able to assign leucine and isoleucine definitively.

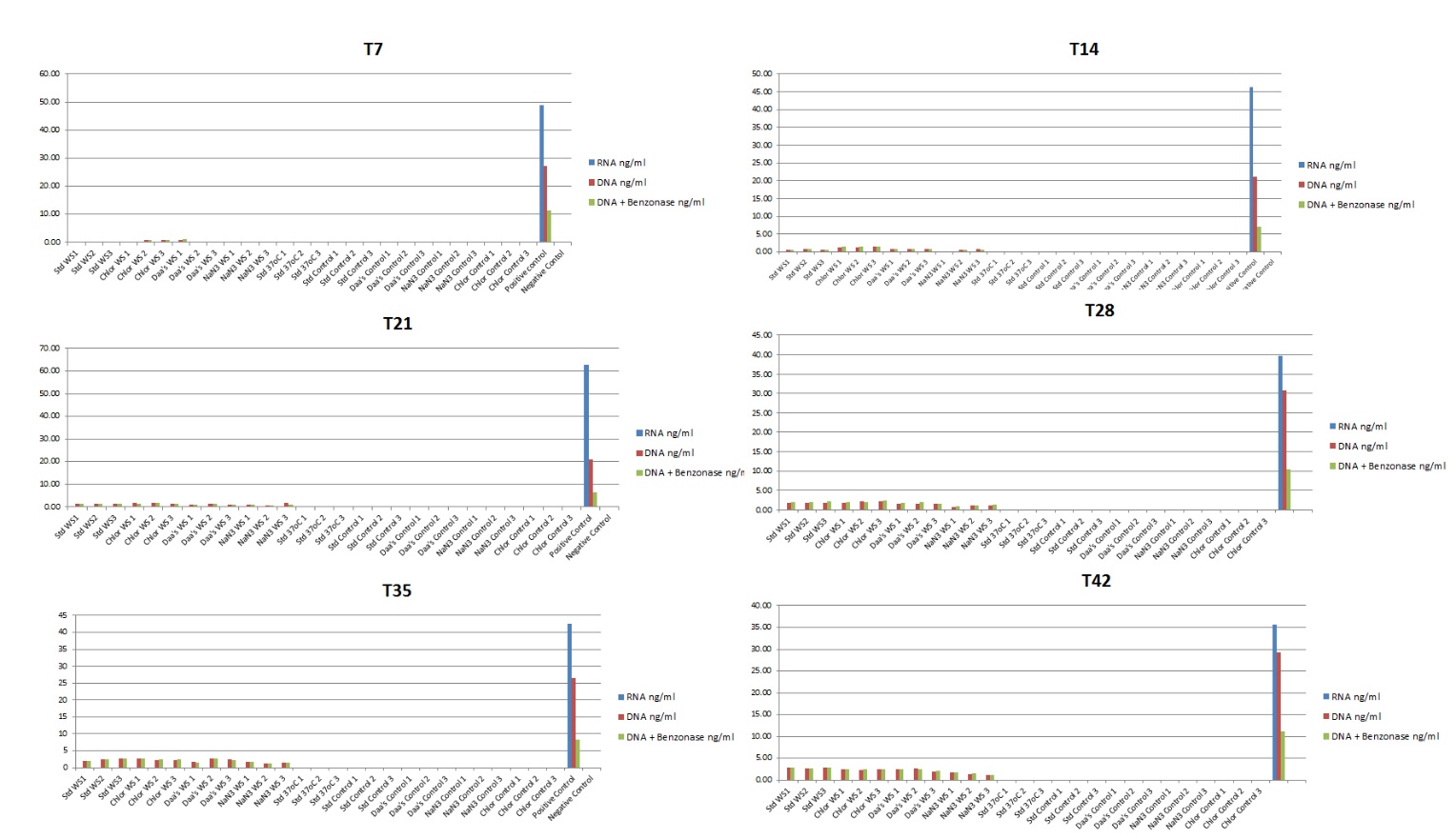
**Figure S2**. RNA and DNA Analysis. All samples were tested for the presence of DNA and RNA using the high sensitivity Qubit kits. No RNA was ever detected, and DNA was detected after the samples changed colour. To verify whether this was in fact DNA, benzonase was applied to the samples and left to incubate. A positive control was always used and the intensity of the positive control always reduced by at least half, while the abio samples “DNA intensity” was never reduced significantly.

**MALDI-TOF Mass Spectrometry of Commercial Amino Acids**

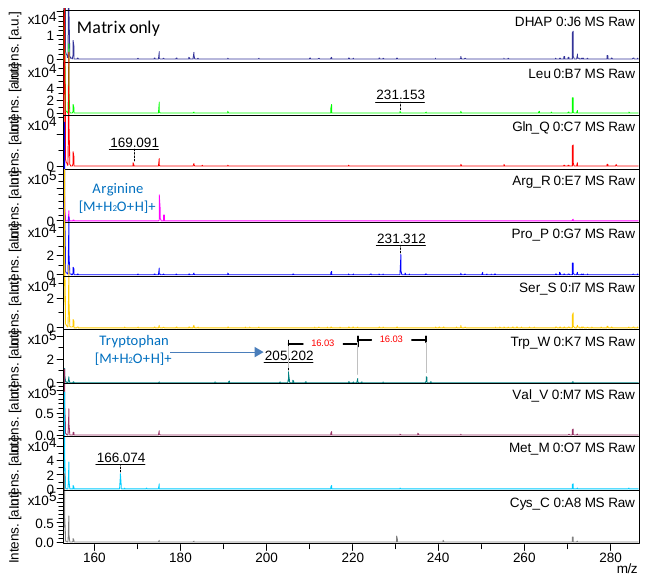
**Figure S3A**. MALDI-TOF spectra of leucine, glutamine, arginine, proline, serine, tryptophan, valine, methionine and cysteine.

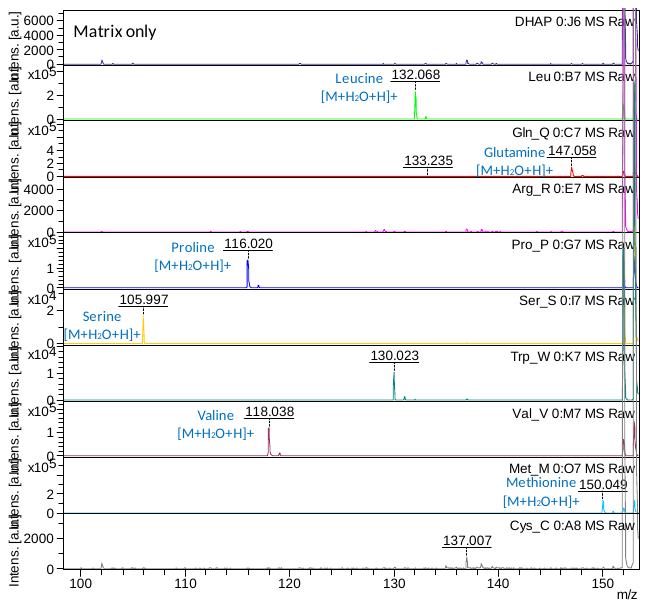
**Figure S3B**. MALDI-TOF zoomed mass spectra of leucine, glutamine, arginine, proline, serine, tryptophan, valine, methionine and cysteine.

**Figure S2C**. MALDI-TOF zoomed mass spectra of leucine, glutamine, arginine, proline, serine, tryptophan, valine, methionine and cysteine.

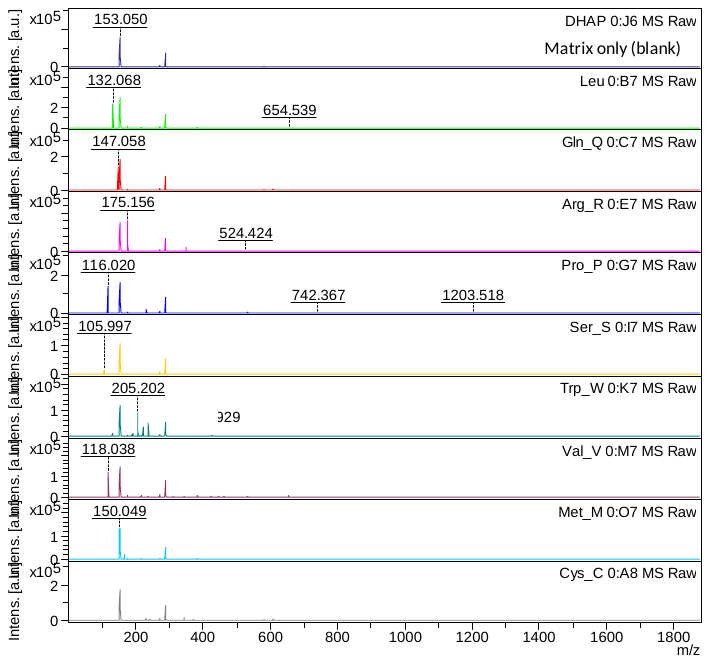

**Figure S3D.** MALDI-TOF mass spectra of tyrosine, asparagine, threonine, phenylalanine, aspartic acid, glycine, isoleucine, lysine, and alanine.

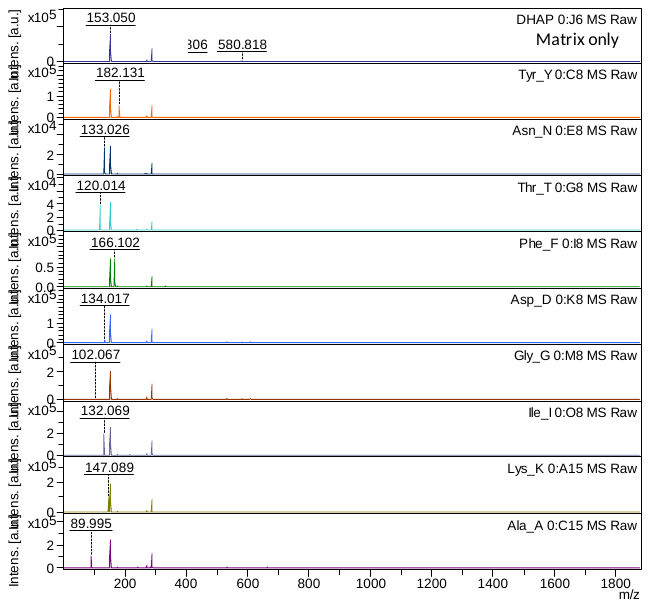

**Figure S3E**. MALDI-TOF zoomed mass spectra of tyrosine, asparagine, threonine, phenylalanine, aspartic acid, glycine, isoleucine, lysine, and alanine.
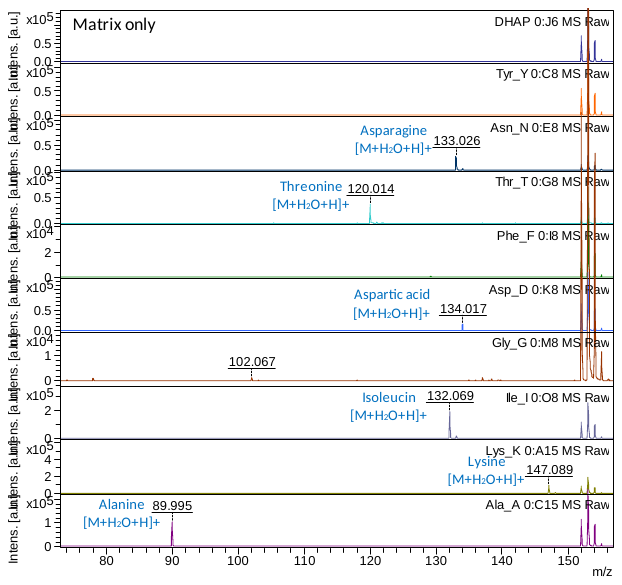

**Figure S3F**. MALDI-TOF zoomed mass spectra of tyrosine, asparagine, threonine, phenylalanine, aspartic acid, glycine, isoleucine, lysine, and alanine.
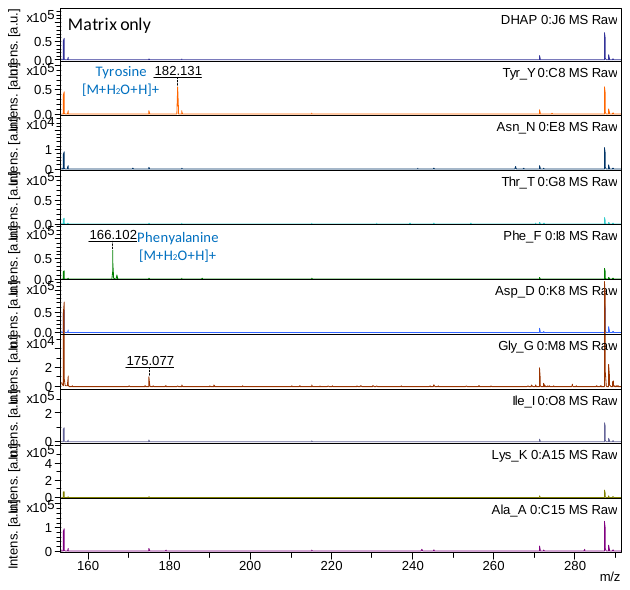

**Figure S3G**. MALDI-TOF mass spectra of glutamic acid and histidine.

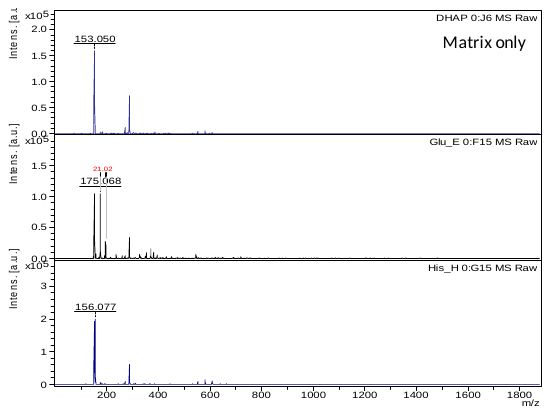

**Figure S3H.** MALDI-TOF zoomed mass spectra of glutamic acid and histidine

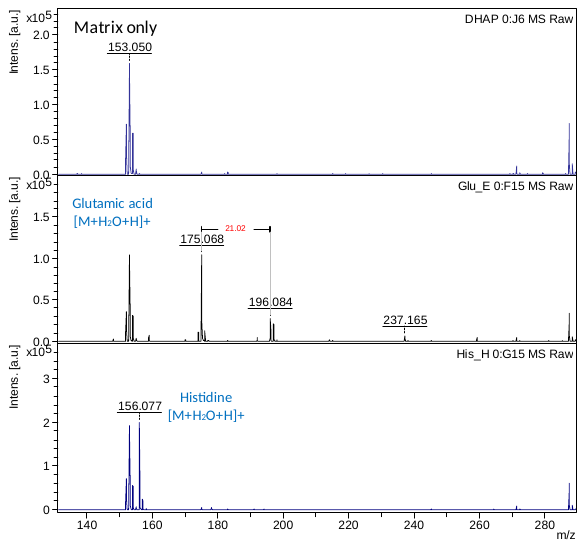

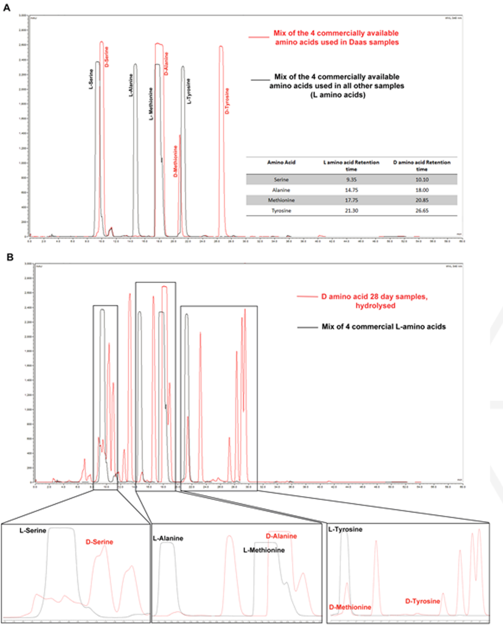

**Figure S4**. Day 28 D amino acid samples were hydrolysed with 6M HCl, and the resulting amino acid mixture was derivatized with Marfeys reagent. The samples were then analysed using reversed phase C18 chromatography to identify if there had been any conversion of D amino acids to L amino acids. There was no detected conversion of D into L, further verifying no presence of bacterial contamination in the samples.

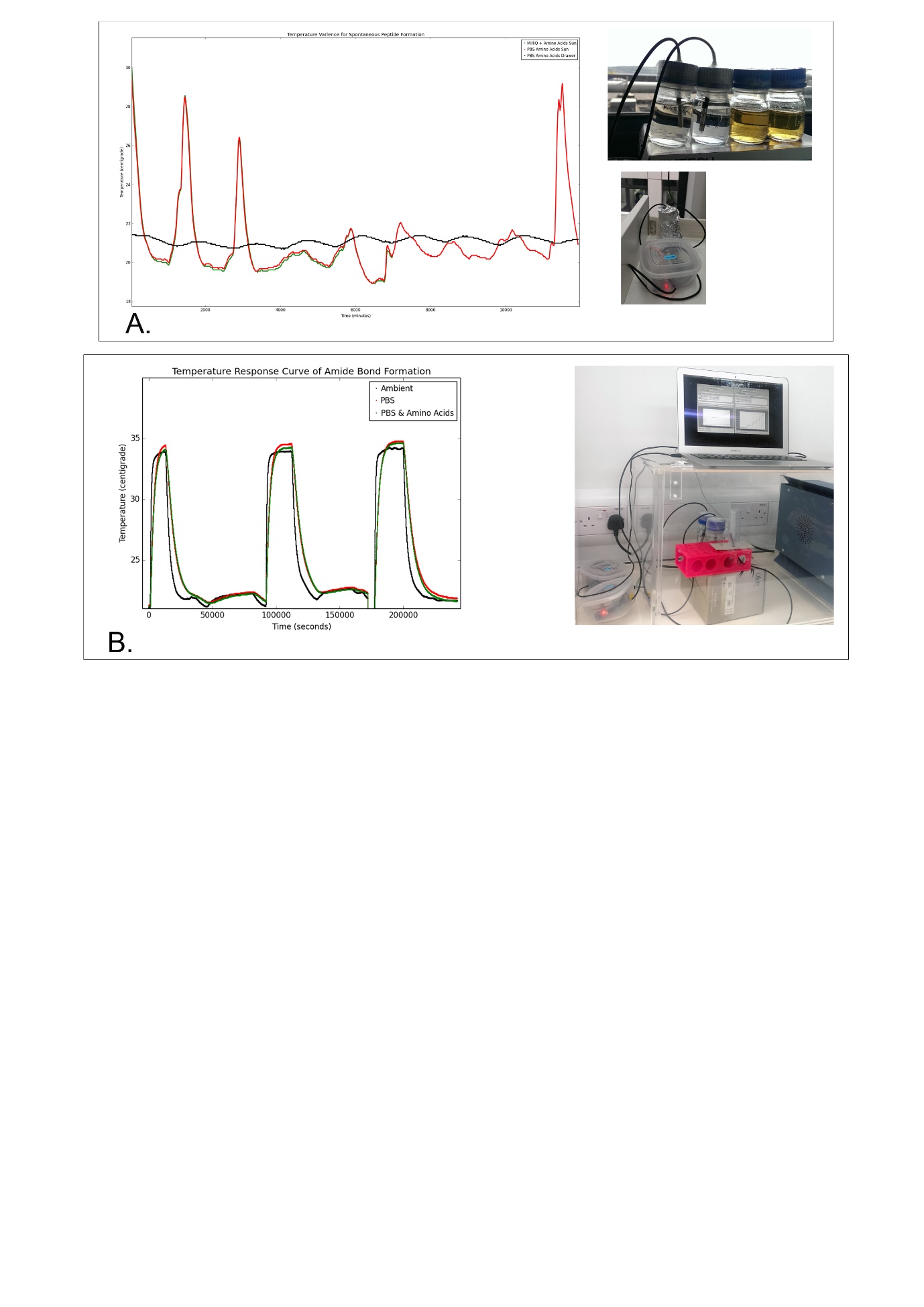
.

**Figure S5**. Bespoke hardware and accompanying software to measure the temperature fluctuations on the windowsill (A), then in a controlled heating environment sans sunlight, to characterize the difference amino acids in solution made, and the amount of energy consumed in the system (B) and finally the completely controlled environment used for all subsequent experiments. All the apparatus were controlled by either Arduino or Raspberry Pi, with Arduino and Python programs respectively. Temperatures were monitored with medical grade temperature sensors.
